# Contrasting transcriptional responses and genetic determinants underlie *Zymoseptoria tritici* adaptation mechanisms to simulated host defense environments

**DOI:** 10.64898/2026.04.14.718361

**Authors:** Silvia Miñana-Posada, Alice Feurtey, Bruce A. McDonald, Cécile Lorrain

## Abstract

Successful colonization of the wheat apoplast requires that *Zymoseptoria tritici* tolerate host-derived stresses, but the mechanisms underlying this adaptation remain poorly understood. We combined phenotypic assays, transcriptomics, and genome-wide association analyses to characterize fungal responses to acidic pH, salicylic acid, gibberellic acid, and oxidative stress. Exposure to salicylic acid inhibited *in vitro* growth across a global collection of 411 *Z. tritici* strains, whereas acidic pH promoted growth, illustrating contrasting effects on pathogen performance of environments simulating host-defense responses. At the transcriptional level, acidic pH and oxidative stress induced the strongest and most similar responses, while salicylic acid elicited a more distinct transcriptional program and gibberellic acid caused only limited transcriptional changes. Although the sets of differentially expressed genes were largely condition specific, overlapping enrichment of transport- and redox-related functions across conditions indicated shared transcriptional responses. K-mer based genome-wide association mapping identified five candidate loci associated with growth under acidic pH, gibberellic acid and salicylic acid, including four loci specific to a single growth condition. These loci colocalized with genes implicated in cell wall remodeling, nitrogen metabolite regulation, proteostasis, and ubiquitin-related processes. This study highlights the multifaceted strategies employed by *Z. tritici* to navigate environments simulating host-defense responses, involving shared and environment-specific adaptations. We provide new insights into the genetic and molecular basis of fungal resilience, with implications for understanding pathogen-host interactions.

## Introduction

The ascomycete fungus *Zymoseptoria tritici*, the causal agent of Septoria tritici blotch (STB), is one of the most damaging wheat pathogens, causing significant global crop losses and threatening food security (Fones and Gurr 2015). Emerging in the Fertile Crescent during the domestication of wheat, *Z. tritici* is thought to have coevolved with its host, developing extensive genetic diversity and phenotypic plasticity (Stukenbrock et al. 2007; Feurtey et al. 2023). This high evolutionary potential is illustrated by the ability of *Z. tritici* populations to rapidly evolve resistance to fungicides and overcome host resistance genes within a few years of their deployment (Plissonneau et al. 2017; Hartmann et al. 2018; Welch et al. 2018). In addition to host-associated selective pressures, adaptation to various abiotic stresses also plays an important role in shaping the evolutionary potential of *Z. tritici* (Dutta et al. 2021). For instance, growth in thermally-stressful environments revealed a significant strain-level variation in temperature tolerance within *Z. tritici* populations (Zhan and McDonald 2011; Zhong et al. 2020; Boixel et al. 2022; Miñana-Posada et al. 2024). Dissecting the genetic architecture underlying such quantitative phenotypic variation remains a major challenge, as traits related to fitness, pathogenicity, and stress tolerance are typically polygenic and governed by complex gene regulatory networks.

Plant defense responses can include the production of phytohormones, reactive oxygen species (ROS), and changes in cellular pH. These defenses are thought to play a critical role in shaping the dynamic interactions between hosts and their pathogens. The pH of various cellular compartments in plant leaf cells is tightly regulated to optimize biochemical functions. For instance, the apoplast and vacuole typically maintain an acidic pH of 5.0 to 6.0, which is essential for ion exchange and signaling, while the cytosol maintains a near-neutral pH for enzymatic activities and metabolic reactions (Tournaire-Roux et al. 2003; Felle et al. 2008; Geilfus 2017). The apoplast pH can change rapidly in response to pathogen infection and other environmental stresses (Geilfus 2017). Phytohormones including salicylic acid (SA) and jasmonic acid (JA) are central to plant immune responses. SA mediates local acquired resistance (LAR) in infected leaves, while JA regulates induced systemic resistance (ISR) (Métraux et al. 2002; Durrant and Dong 2004). These pathways often interact antagonistically to fine-tune defense responses (Spoel et al. 2007; Pieterse et al. 2009). Other phytohormones, like gibberellic acid (GA), are primarily involved in growth and development (Gupta and Chakrabarty 2013), but their role in plant defense is poorly understood. Some studies suggest that GA may reduce pathogen virulence (Eshel et al. 2002; Tanaka et al. 2006), while others report no significant effects on pathogen growth and disease development (Buhrow et al. 2021). GA is produced by some fungal species and can play a key role in the infection process as described for the rice pathogen *Fusarium fujikuroi* (Desjardins et al., 2000; Zhang et al., 2019). Oxidative stress, marked by production of ROS during an oxidative burst, is an early cellular response to pathogen detection that influences host signaling and pathogen virulence (Torres et al. 2006). With the exception of one study using ROS (Zhong et al. 2021), no studies have investigated the adaptive mechanisms underlying the direct effects of these factors related to plant defense on *Z. tritici*.

Host-produced phytohormones and ROS can act as direct chemical stressors and affect fungal physiology. Multiple plant hormones were shown to have specific effects on fungal processes, including alterations to growth and morphogenesis (e.g., auxin-induced invasive vs pseudo-hyphal growth)(Rao et al. 2010), transcriptional rewiring of core growth pathways (e.g., abscisic acid effects on cell-cycle gene expression)(Xu et al. 2018), disruption of membrane homeostasis (e.g., salicylic acid-induced inhibition of ergosterol biosynthesis and reduced membrane integrity)(Kong et al. 2021), and interference with trafficking and cytoskeletal dynamics (e.g., cytokinin-driven perturbation of endocytosis/trafficking in plant pathogens)(Gupta et al. 2021). ROS produced during the plant oxidative burst imposes oxidative damage such as lipid peroxidation, protein oxidation, and DNA lesions, but also functions as a signal that activates conserved fungal stress-response circuits (Singh et al. 2021). Plant-associated fungi typically counter ROS via inducible antioxidant and redox-buffering systems including superoxide dismutases, catalases, peroxidases/peroxiredoxins, and glutathione- and thioredoxin-dependent pathways (Park and Son 2024). These pathways are coordinated by regulators such as the AP-1-like bZIP transcription factor Yap1, the two-component response regulator Skn7, and MAPK pathways including Hog1, which collectively control detoxification, repair, and broader metabolic adaptation (Lev et al. 2005; Fassler & West 2011; Simaan et al. 2019; Park & Son 2024). Importantly, this oxidative stress-response network often intersects with virulence-associated functions in plant pathogens (e.g., transport, iron homeostasis, and development), meaning that phytohormones and ROS exposure can jointly shape quantitative variation in growth and stress tolerance traits that are relevant to infection (Chen et al. 2014; Chen et al. 2017; Qi et al. 2019; Vangalis et al. 2021).

Here we employ a multi-omics approach to investigate the molecular and phenotypic mechanisms by which *Z. tritici* adapts to different environments that may be associated with host-defense responses. By integrating phenotypic assays, transcriptomics, and genome-wide association studies (GWAS), we aim to uncover how pH, phytohormones and oxidative stress influence fungal growth. Using a global collection of 411 *Z. tritici* strains, we evaluated *in vitro* phenotypic traits under five growth conditions. Our approach leverages multi-reference k-mer-based GWAS to identify genetic loci associated with particular phenotypic traits, complemented by transcriptomic profiling to characterize differential gene expression patterns across treatments. We hypothesized that (I) exposure to different environments associated with host-defense responses impacts fungal growth, reflected by changes in key phenotypic traits; (II) phenotypic responses to distinct environments simulating host defenses are associated with specific genomic loci; and (III) differential gene expression patterns vary, revealing distinct yet potentially overlapping pathways activated by environments simulating host defense responses. This study seeks to unravel the main mechanisms driving *Z. tritici*’s adaptive responses to different components of the local environment that is produced as part of a plant’s defense-response.

## Results

### Growth dynamics and sensitivity in four environments simulating host defense responses in 411 *Z. tritici* strains

To assess the effects of environments simulating host defense responses on *Z. tritici*, we phenotyped 411 strains from eight genetic clusters (i.e. populations; Fig. 1A; Supplementary Table S1; (Miñana-Posada et al. 2024). Seven *in vitro* growth variables were measured under five environments: neutral pH (GPL pH 7), acidic pH (GPL pH 5), hydrogen peroxide (H_2_O_2_, pH 7), gibberellic acid (GA, pH 5), and salicylic acid (SA, pH 5). We observed interdependence among growth variables, with significant correlations between pairs of growth variables observed in 78% of comparisons (466 out of 595; Fig. 1B; Supplementary Fig. S1), with Pearson’s r values ranging from -0.61 to 0.99 (Benjamini-Hochberg (BH) adjusted p-value < 0.05; Supplementary Table S2; Supplementary Fig. S1). The empirical area under the curve (eAUC) was consistently and positively correlated with carrying capacity and spore concentrations across environments (Fig. 1B), indicating that eAUC effectively reflects overall pathogen growth (carrying capacity: r = 0.71–0.93; concentrations: r = 0.57–0.99, all BH-adjusted p-value < 0.05). Growth rate exhibited a consistent negative correlation with inflection time within environments (r = -0.61 to -0.25, all BH-adjusted p-value < 0.05), but only weak correlations with carrying capacity (r = -0.10 to 0.17; significant only in GPL pH 7 and H_₂_O_₂_ pH 7; BH-adjusted p-value < 0.05). Growth rate correlations varied between environments, with significant positive correlations primarily among GPL pH 7, GPL pH 5, GA pH 5, and H_₂_O_₂_ pH 7 (r = 0.21–0.61), whereas growth under SA showed no significant correlation with growth rate in other environments. In contrast, inflection times were positively correlated across all environments (r = 0.20–0.63, all BH-adjusted p < 0.05). Together, these results suggest that *Z. tritici* strains show consistent performance across environments simulating host defense responses for carrying capacity, eAUC, and spore concentrations, whereas growth rate is more environment dependent.

**Fig. 1.**
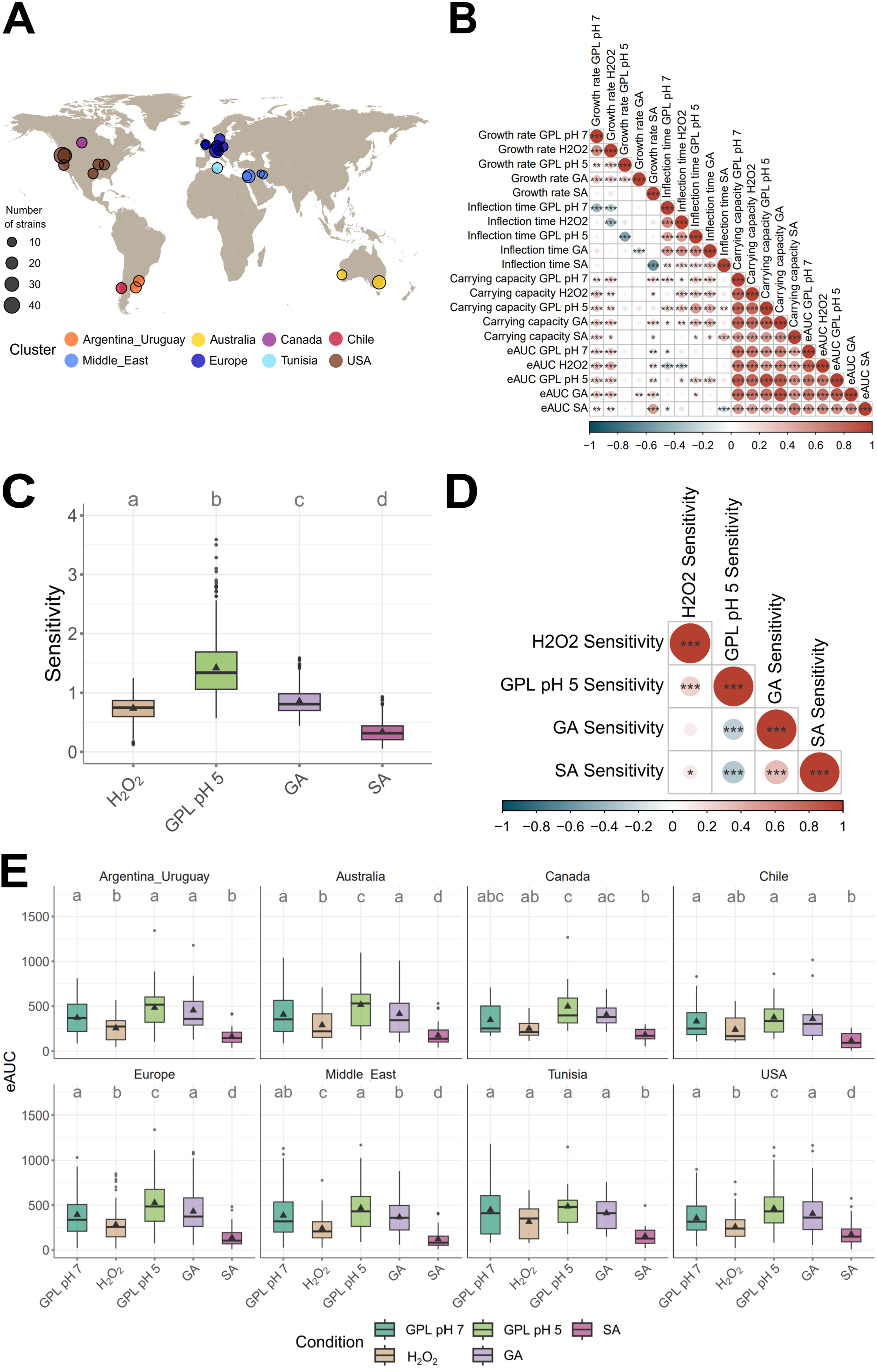
*In vitro* phenotypic response of 411 *Zymoseptoria tritici* strains to four environments simulating host defense responses. (A) Geographic distribution and genetic cluster assignment of 411 *Zymoseptoria tritici* strains. Circle sizes indicate the number of strains from each sampling location, with only locations containing more than five strains shown. Colors represent distinct genetic clusters as defined by Miñana-Posada et al. (2024). (B) Pearson’s correlations between growth rate, inflection time, carrying capacity, and eAUC under simulated plant defense responses. The color scale represents the correlation coefficients, while significance levels are denoted by asterisks (*: Benjamini–Hochberg corrected p-value < 0.05; **: Benjamini–Hochberg corrected p-value < 0.01; ***: Benjamini–Hochberg corrected p-value < 0.001). (C) Boxplots illustrating per strain sensitivity as ratios between treatments and control media conditions. These include neutral pH treatments (15 hpi and 90 hpi H_₂_O_₂_) or low pH media (GPL pH 5) compared to neutral control media (GPL pH 7), and low pH treatments (GA, SA) compared to low pH media (GPL pH 5). Triangles within each boxplot indicate the mean sensitivity for each condition. Lowercase letters (a–d) represent significant differences between treatments, determined by Tukey’s post-hoc pairwise comparisons based on a linear mixed-effects model. (D) Pearson’s correlations of sensitivities across environments simulating host-defense responses. The color scale and significance levels follow the conventions of panel (B). (E) Boxplots showing per strain eAUC, grouped by genetic cluster. Triangles within each boxplot indicate the mean sensitivity for each condition and genetic cluster. Lowercase letters (a–e) indicate significant differences between environments simulating host-defense responses within genetic clusters, as assessed by Tukey’s post-hoc pairwise comparisons based on a linear mixed-effects model.

We observed significant differences in the growth variable variance across environments (Levene’s test, p-value < 0.001), suggesting distinct phenotypic impacts according to environment. Linear mixed-effects models (LMERs) confirmed significant environment effects on all growth variables (ANOVA p-value < 0.001; Supplementary Table S3). Post hoc analyses showed that environments differed significantly with few exceptions, such as eAUC GPL pH 7 and eAUC GA pH 5 (Tukey p-value > 0.05; Supplementary Fig. S2A; Supplementary Table S4). Ranking of mean eAUC values across environments, from lowest to highest, revealed the following order: SA pH 5, H_2_O_2_ pH 7, GPL pH 7, GA pH 7, and GPL pH 5 (Supplementary Table S5). For growth rate, the environments, from lowest to highest, ranked as follows: SA pH 5, GPL pH 7, H_2_O_2_ pH 7, GA pH 5, and GPL pH 5. We found a significant difference for growth rates between GPL pH 5 vs SA pH 5 (Tukey p-value < 0.05). However, other pairwise comparisons showed no significant differences, specifically between GPL pH 7 vs H_2_O_2_ pH 7, GA pH 5 vs H_2_O_2_ pH 7, and GPL pH 5 vs GA pH 5 (Tukey p-value > 0.05; Supplementary Fig. S2B; Supplementary Table S4).

To further evaluate environment-specific effects on growth, we calculated sensitivity as eAUC ratios relative to baseline environments (GPL pH 7 for neutral conditions or GPL pH 5 for acidic conditions, i.e. the SA and GA conditions; Fig. 1C). Ratios below 1 indicated reduced growth (i.e., sensitivity), while ratios above 1 signified enhanced growth. In the absence of any other factor, acidic pH consistently promoted growth compared to neutral pH. Environments with H_2_O_2_, GA, and SA significantly suppressed growth (Fig. 1C). The SA environment elicited the strongest growth reduction compared to the control environment with the corresponding pH (pH 7 and pH 5, respectively; Supplementary Table S4-S5). Correlation analyses revealed distinct patterns of sensitivity relationships among environments. Positive correlations were observed among H_2_O_2_, and GPL pH 5 sensitivities. Conversely, sensitivity to GA and SA negatively correlated with sensitivity to GPL pH 5, indicating that strains most sensitive to GA or SA tended to thrive under acidic environments (Fig. 1D; Supplementary Table S2). These findings demonstrate that GPL pH 5 enhances the growth of *Z. tritici*. In contrast, salicylic acid exerts strong inhibitory effects, highlighting the diverse phenotypic responses of this pathogen to environments simulating host defense responses.

Finally, we examined whether the environments simulating host defense responses had distinct effects on sensitivity, eAUC, and growth rate across strains from different genetic clusters of *Z. tritici* populations. No significant differences in sensitivity were detected between genetic clusters within the same environment (Tukey p-value > 0.05; Supplementary Fig. S3; Supplementary Table S6). Similarly, eAUC values did not vary significantly between clusters within each environment (Tukey p-value > 0.05; Supplementary Fig. S4; Supplementary Table S6). However, comparisons of environments per genetic cluster revealed distinct patterns of significance, demonstrating that the relationships between environments are not uniform across genetic clusters (Fig. 1E; Supplementary Fig. S4; Supplementary Fig. S5; Supplementary Table S6; Supplementary Fig. S3B).

### Environments simulating host-defense responses elicit specific transcriptional programs, with convergence between acidic pH and oxidative stress responses

We examined the transcriptomic responses of the *Z. tritici* IPO323 reference strain under the same five environments used in phenotyping: GPL pH 7, GPL pH 5, salicylic acid (SA), gibberellic acid (GA), and hydrogen peroxide (H_2_O_2_). To take into account the changes across growth stages, transcriptomic profiles were analyzed after 15 hours for the control condition (15 hpi GPL pH 7) and H_2_O_2_ (15 hpi H_2_O_2_) and at 90 hours for GPL pH 7, GPL pH 5, H_2_O_2_ pH 7, SA pH 5 and GA pH 5 (Fig. 2A). A principal component analysis (PCA) of normalized read counts revealed distinct clusters corresponding to the replicates from the same environments, reflecting differences in gene expression profiles induced by environments simulating host-defense responses (Fig. 2B). Along PC1 (42% of the variance), the 15 hpi control (15 hpi GPL pH 7) and 15 hpi H_2_O_2_ samples clustered closely together, whereas the 90 hpi control (GPL pH 7) samples formed a distinct cluster. The 90 hpi H_2_O_2_ samples were displaced from the 15 hpi H_2_O_2_ samples and overlapped with the GPL pH 5 environment. Along PC2 (35% of the variance), samples grown in GPL at acidic pH (GPL pH 5) and at neutral pH (GPL pH 7) show clear separation, implying a significant transcriptional shift under acidic environments. Samples grown under the salicylic and gibberellic acid environments also formed discrete clusters that were distinct from the acidic pH media (GPL pH 5; Fig. 2B). Overall, these results show that IPO323 displays distinctive transcriptomic responses to the different environments simulating host-defense responses used in our experiments, suggesting that different regulatory pathways were triggered by these environments.

**Fig. 2.**
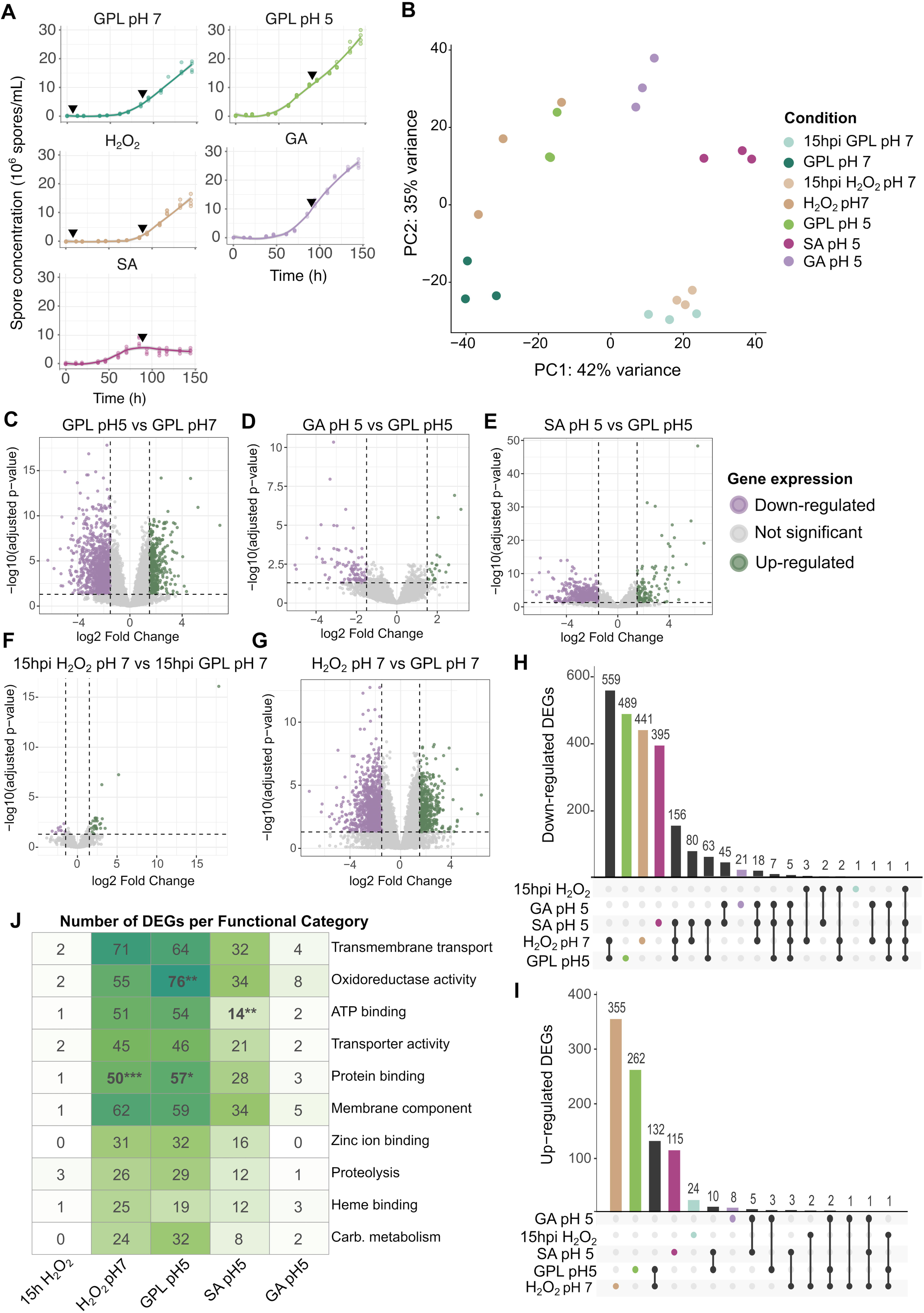
Transcriptome responses of *Zymoseptoria tritici* to four environments simulating host-defense responses. A) Growth curves of the reference strain IPO323 under the neutral media control (GPL pH 7) and in four environments simulating plant defense responses: acidic media control (GPL pH 5); hydrogen peroxide (H_2_O_2_); gibberellic acid pH 5 (GA); salicylic acid pH 5 (SA). Points represent replicate measurements for each timepoint and treatment, while lines indicate locally estimated scatterplot smoothing (LOESS) regressions. Black triangles mark the timepoints used for transcriptomic analyses for each condition (i.e., 15 or 90 hours post-inoculation). B) Principal component analysis (PCA) of transcriptomes from all environments and timepoints for the reference strain IPO323, displaying the first two principal components. Points represent the three replicates per environment and timepoint, with colors indicating different treatments and timepoints. C-G) Volcano plots showing differential expression for each environment relative to its corresponding control: C) GPL pH 5 vs GPL pH 7, D) GA pH 5 vs GPL pH 5, E) SA pH 5 vs GPL pH 5, F) 15 hpi H_2_O_2_ pH 7 vs 15 hpi GPL pH 7, G) 90 hpi H_2_O_2_ pH 7 vs 90 hpi GPL pH 7. Vertical dashed lines indicate the log2 fold-change thresholds (±1.5) and the horizontal dashed line notes the DESeq2 adjusted p-value cutoff (p-adjusted < 0.05). Points are colored by expression status (up-regulated in green, down-regulated in purple, or not significant in grey). H–I) Shared and specific differentially expressed genes (DEGs) across environments for (H) down-regulated and (I) up-regulated gene sets. Bars report the number of DEGs in each intersection, and the dot-line matrix below indicates the corresponding combination of environments contributing to each intersection. Treatment-specific DEGs are highlighted by colors. J) Distribution of annotated DEGs across broad functional categories for each environment. Cells report the number of DEGs assigned to each category, and asterisks indicate functional categories significantly enriched within a treatment relative to the set of functionally annotated genes in the IPO323 genome (two-side Fisher’s exact test with FDR correction; * p-value < 0.05, ** p-value < 0.01, *** p-value < 0.001).

We identified differentially expressed genes (DEGs) by comparing each environment to their respective controls under neutral or acidic pH environments (Fig. 2C-G; Supplementary Table S7). A total of 4,517 DEGs (33.7% of IPO323 genes) were detected across environments, defined by a log2-fold change greater than 1.5 or less than -1.5 and an adjusted p-value of < 0.05 (Supplementary Table S7). Among these, 1,088 DEGs (24.1% of the total DEGs) were upregulated in at least one environment compared to its corresponding control, while the majority of DEGs (75.9%, 3,429) were downregulated (Fig. 2C-G). The most extensive responses occurred for the 90 hpi oxidative stress and acidic pH environments which triggered broad changes in gene expression. The 90 hpi H_2_O_2_ (90 hpi) environment induced 1,776 DEGs (502 upregulated and 1,274 downregulated), and the GPL pH 5 environment induced a comparable 1,694 DEGs (411 upregulated and 1,283 downregulated; Fig. 2C & 2G). This indicates that low pH alone elicits a transcriptomic shift on a scale that is similar to that induced by oxidative stress (Fig. 2B). Salicylic acid produced an intermediate response (905 DEGs; 134 upregulated and 771 downregulated), consistent with a pronounced but more specific remodeling of gene expression relative to the pH 5-driven baseline (Fig. 2E). In contrast, gibberellic acid had only a minor effect (118 DEGs; 20 upregulated and 98 downregulated), suggesting limited perturbation of IPO323 transcription relative to the pH 5-driven baseline (Fig. 2D). Similarly, the early oxidative stress response was minimal. At 15 hpi, H_2_O_2_ triggered only 36 DEGs (27 upregulated and 9 downregulated), aligning with the close clustering of 15 hpi GPL pH 7 and 15 hpi H_2_O_2_ samples. This pattern indicates that large-scale transcriptional rewiring emerges primarily following prolonged exposure and/or at later growth stages (Fig. 2F-G). Despite this limited response, the 15 hpi H_2_O_2_ treatment induced the highest upregulated gene observed across all environments and time points: ZtIPO323_085730 (encoding a protein of unknown function), with log2FC = 17.8 at 15 hpi H_2_O_2_ and log2FC = 3.8 at 90 hpi H_2_O_2_ relative to the respective control (Fig. 2F-G; Supplementary Table S7). In contrast, ZtIPO323_085730 was not significantly differentially expressed in GPL pH 5 vs GPL pH 7 nor in GA vs pH 5, and it was downregulated in SA vs pH 5 (log2FC = -6.3) (Supplementary Table S7).

To determine whether these responses converge on common pathways, we next examined the overlap of DEGs across environments to identify genes that were regulated similarly across different environments simulating host-defense responses. The overlap between environments was limited (Jaccard range 0.005-0.335), consistent with most DEGs being environment-specific. The GPL pH 5 and 90 hpi H_2_O_2_ environments showed the highest similarity (Jaccard = 0.335) and the largest shared set of DEGs with 559 shared downregulated DEGs and 132 shared upregulated DEGs (Fig. 2H-I; Supplementary Fig. S6). Outside of this pair, overlaps were very limited. SA showed partial similarity with both 90 hpi H_2_O_2_ (Jaccard = 0.164) and GPL pH 5 (Jaccard = 0.129). The 15 hpi H_2_O_2_ environment showed almost no overlap with the other environments (Jaccard = 0.005-0.016; Supplementary Fig. S6A). In total, only nine DEGs were shared across four out of five environments (GPL pH5 vs GPL pH7, SA vs GPL pH5, GA vs GPL pH5, 90 hpi H_2_O_2_ vs 90 hpi GPL pH 7, and 15 hpi H_2_O_2_ vs 15 hpi GPL pH 7; Supplementary Fig. S6A-B). Six of these shared DEGs were consistently downregulated across comparisons, including five genes of unknown function (ZtIPO323_025910, ZtIPO323_059230, ZtIPO323_118340, ZtIPO323_038710, and ZtIPO323_077710) and one gene coding for a predicted glycoside hydrolase family 16 (ZtIPO323_089860). Three shared genes displayed environment-dependent expression, including ZtIPO323_068440 (unknown function) which was induced under oxidative stress and low pH (up to 2.91 log2FC) but repressed under SA (-2.41 log2FC), ZtIPO323_088430 (alcohol oxidase) which was strongly induced in 90 hpi H_2_O_2_ (2.86 log2FC) but downregulated in GA, SA, and acidic pH (down to -4.98 log2FC), and ZtIPO323_124290 (a predicted effector) that was induced in GA (1.71 log2FC) and SA (1.72 log2FC) but repressed in GPL pH 5 (-2.23 log2FC) and 90 hpi H_2_O_2_ (-2.30 log2FC)(Supplementary Fig. S6C). Together, these results show that transcriptional responses to environments simulating host-defense responses are predominantly environment-specific, with only limited overlap across environments apart from acidic pH and 90 hpi oxidative stress.

We next asked whether the different environments simulating host-defense responses converged on similar functional responses by comparing the distribution of annotated DEGs across broad functional categories (Fig. 2J). Although many DEGs lacked functional annotation, the functionally annotated subset associated to membrane-associated processes and transport (e.g., transmembrane transport, transporter activity, membrane component) as well as redox-associated functions (oxidoreductase activity), suggesting that there may be conservation at the level of biological function even when the underlying DEG sets differ. The *Z. tritici* response to the GPL pH 5 environment consistently showed significant enrichment for oxidoreductase activity (Fisher’s exact test, FDR-adjusted p-value < 0.05; Fig. 2J), whereas the H_2_O_2_ (90 hpi) environment was significantly enriched for protein binding (Fisher’s exact test, FDR-adjusted p-value < 0.05; Fig. 2J). Finally, salicylic acid treatment showed enrichment for ATP binding (Fisher’s exact test, FDR-adjusted p-value < 0.05; Fig. 2J). Together, these results indicate that *Z. tritici* responses to environments simulating host-defense responses involve broad functional stress programs, particularly associated with transport and redox-related processes, yet show substantial divergence in the specific genes contributing to each response.

We therefore focused on environment-specific transcriptional regulation, as specific responses accounted for the majority of both repressed and induced genes (Fig. 2H–I). We identified 489 pH 5-specific down-regulated and 262 up-regulated DEGs (GPL pH 5 vs GPL pH 7), with genes encoding for oxidoreductase activity significantly enriched among the up-regulated genes (Fisher’s exact test, FDR-adjusted p-value < 0.05; Supplementary Fig. S7-S8). Oxidative stress (90 hpi H_2_O_2_) elicited 441 specific down-regulated DEGs, which were enriched for protein binding functions (Fisher’s exact test, FDR-adjusted p-value < 0.05), and 355 up-regulated DEGs that showed no significant functional enrichment (Fig. 2H–I; Supplementary Fig. S7). The salicylic acid environment resulted in 395 SA-specific down-regulated and 115 up-regulated DEGs without significant enrichment for any function (Fig. 2H–I; Supplementary Fig. S7). Interestingly, SA markedly induced a salicylate hydroxylase (ZtIPO323_037930: 6.2 log2FC, p-adjusted < 0.05) and a candidate effector gene (ZtIPO323_019470: 6.7 log2FC, p-adjusted < 0.05), indicating activation of an SA-catabolic/detoxification program alongside putative virulence-associated transcripts (Supplementary Fig. S8). In contrast, GA and 15 hpi H_2_O_2_ produced few environment-specific DEGs, with 21 down-regulated and 8 up-regulated DEGs for GA, and 1 down-regulated and 24 up-regulated DEGs for 15 hpi H_2_O_2_, including three genes annotated with proteolytic functions (Supplementary Fig. S7). Taken together, these results indicate that, with the exception of GA, the tested environments simulating host-defense responses elicit specific transcriptional responses characterized by only nine shared DEGs and from 44% (GPL pH 5) to 56% (SA) environment-specific DEGs.

### K-mer-based GWAS uncovers loci associated with adaptation to low pH and salicylic acid

To elucidate the genetic factors affecting growth in environments simulating host-defense responses, we performed a k-mer-based genome-wide association study (GWAS) on 411 strains. Our analysis evaluated seven phenotypic traits (i.e., carrying capacity, eAUC, growth rate, inflection and spore concentration 48h-, 96h- and 144hpi) in five growth environments (i.e., GPL pH 7, GPL pH 5, salicylic acid, gibberellic acid and hydrogen peroxide), and the trait sensitivity under the four non-control environments (i.e. relative growth trait compared to GPL pH 7). This k-mer-based GWAS identified 191 significant k-mers associated with 19 phenotypes, applying a 10% family-wise error rate threshold (Supplementary Table S8). We mapped the significant k-mers to eight genomic loci based on their proximity to orthologous genes. After excluding loci overlapping with transposable elements (TEs), five loci were retained, each containing more than two significant k-mers (Supplementary Table S9).

We identified one locus on chromosome 3 associated with eAUC in the salicylic acid environment, represented by 33 significant k-mers within a conserved genomic region present in all 19 reference genomes (Fig. 3; Supplementary Table S9). This locus is located 24 bp upstream of orthogroup OG_5933, which encodes a predicted beta-1,3-glucanase (ZtIPO323_043860; Supplementary Table S9). In the reference strain STCH99_1A5, however, the same locus lies 12.4 kbp upstream of OG_5933 due to a large retrotransposon insertion, suggesting transposable element–mediated structural variation potentially affecting this salicylic acid stress-associated gene in STCH99_1A5 (Fig. 3C; Supplementary Table S9). In IPO323, ZtIPO323_043860 was not significantly differentially expressed in the tested environments (Supplementary Table S7). A second locus associated with carrying capacity in the salicylic acid environment was identified on chromosome 8 (Supplementary Table S9). This locus comprised 3 to 6 significant k-mers across all 19 reference genomes. In 13 references, it was located 491-597 bp downstream of orthogroup OG_11569, whereas in the remaining six, it was found in intergenic regions approximately 2 kb from orthogroup OG_7548 (Supplementary Table S10). Manual inspection of sequence alignments, however, confirmed that both orthogroups correspond to ZtIPO323_095260 in the IPO323 reference genome, which encodes a predicted nitrogen metabolite repression protein, NMR-like. The inconsistent ortholog annotations likely result from gene prediction errors rather than true biological differences (Supplementary Table S9). ZtIPO323_095260 was found slightly down-regulated in the 90 hpi H_2_O_2_ vs GPL pH 7 treatment, which suggests a function related to broad responses to environments simulating host-defenses (Supplementary Table S7).

**Fig. 3.**
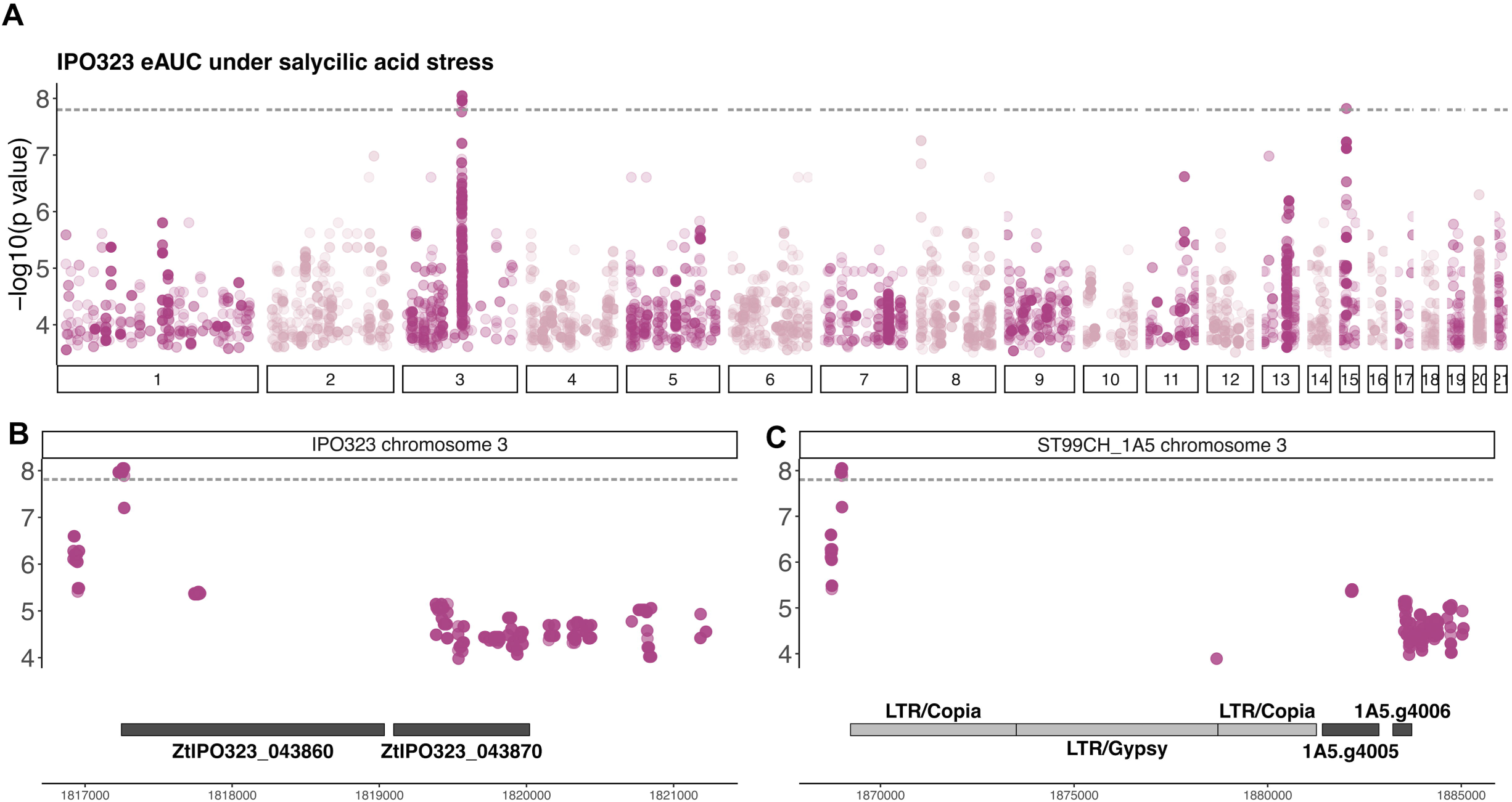
Genetic architecture of salicylic acid adaptation in *Zymoseptoria tritici*. A) Manhattan plot showing k-mer GWAS results for the eAUC under salicylic acid (SA) environment, aligned with the IPO323 reference. The significant k-mer associations are marked strictly above the 10% family-wise error rate significance threshold. (B) Manhattan plot displaying a ∼4.5 kb region on chromosome 3 with significant k-mers associated with eAUC under the SA environment, aligned with the IPO323 reference and located upstream of a predicted beta-1,3-glucanase gene (ZtIPO323_043860). (C) Manhattan plot of a ∼17 kb region on chromosome 3, presenting significant k-mers associated with eAUC under the SA treatment, aligned with the ST99CH_1A5 reference and situated upstream of transposable elements (TEs), followed by a predicted beta-1,3-glucanase gene (1A5.g4005). Horizontal lines indicate the 10% family-wise error rate significance threshold.

Three loci were identified in association with the growth phenotypes found in the gibberellic acid environment. During the early growth stage (48 hpi), one locus on chromosome 11 was identified, encompassing 3–6 significant k-mers across all reference genomes. This locus overlapped with orthogroup OG_10681 in 13 references and was located 462 bp upstream of another orthogroup, OG_2407, in six references (Supplementary Table S9). As observed previously, manual inspection of the two orthogroups confirmed that they correspond to the same gene, annotated as ZtIPO323_114180 in the reference IPO323. The latter encodes an ATP-dependent protease with functions linked to posttranslational modifications, protein turnover, and molecular chaperones but was not significantly differentially expressed in IPO323 (Supplementary table S7). Finally, two loci were identified at the late growth stage (144 hpi). The first locus, located on chromosome 3, contained three significant k-mers in 16 reference genomes, overlapping with orthogroup OG_6945, which encodes a small secreted protein potentially involved in adaptation to environments simulating host-defense responses (ZtIPO323_039430 in IPO323, not differentially expressed). The second locus on chromosome 8 comprised three significant k-mers in 18 references and 30 significant k-mers in the UR95 reference genome. This locus is colocalized with orthogroup OG_7321, which encodes an E3 ubiquitin ligase-like protein (ZtIPO323_092450 in IPO323, not differentially expressed), a gene associated with posttranslational modification processes. Interestingly, this locus was also significantly associated with the GPL pH 5 environment (Supplementary Table S9).

## Discussion

Plant-pathogen interactions are shaped by complex defense mechanisms employed by the host, including phytohormones, reactive oxygen species, and pH regulation. In this study, we investigated the responses of *Z. tritici* to environments simulating host defense responses, including exposure to two phytohormones, oxidative stress, and an apoplast-like pH. By measuring growth dynamics in more than 400 global strains, analyzing differential gene expression patterns in the reference strain, and conducting k-mer based genome-wide association analyses in more than 400 global strains, we provide a comprehensive overview of fungal adaptation to these conditions. Our results show that *Z. tritici* growth is strongly constrained by salicylic acid, though acidic pH by itself promotes fungal growth, underscoring the contrasting effects of different environments the pathogen is likely to experience while infecting its host. More broadly, adaptation of *Z. tritici* to simulated host environments appears to operate across three interconnected levels: phenotypic response, environment-specific transcriptional reprogramming, and underlying genomic variation. At the phenotypic level, salicylic acid was the strongest inhibitor of growth, while acidic pH enhanced performance, indicating that host-associated conditions can either restrict or favor pathogen development depending on the nature of the host environment. At the transcriptional level, acidic pH and oxidative stress produced the broadest and most overlapping responses, consistent with partial convergence on a shared transcriptome program. By contrast salicylic acid induced a more specific regulatory response, while gibberellic acid had little effect on transcription. At the genomic level, k-mer-based GWAS identified a relatively small number of loci associated with growth variation under salicylic acid, gibberellic acid, and acidic pH, pointing to candidate genes involved in cell wall remodeling, nitrogen metabolite regulation, proteostasis, and ubiquitin-related functions. Collectively, these results provide an integrated multi-omics study to better understand how *Z. tritici* responds to the wheat apoplast environment expected following activation of host defenses.

Among the tested environments simulating host-defense responses, we observed that salicylic acid consistently inhibited fungal growth across all strains in our study, highlighting its effectiveness in constraining fungal pathogen growth (Qi et al. 2012). This result is consistent with previous studies demonstrating that SA suppresses colony growth and virulence in diverse fungal species, including *Z. tritici*, *Magnaporthe oryzae*, *Botrytis cinerea*, and *Fusarium spp.* (Mandal et al. 2009; Qi et al. 2012; Dieryckx et al. 2015; Mejri et al. 2019; Yang et al. 2019). The widespread and consistent efficacy of SA across our global strain collection suggests a conserved mechanism underlying its antifungal effects. Our transcriptomic analyses support this hypothesis, as they revealed an SA-specific response in *Z. tritici* involving genes linked to metabolism and cell wall remodeling. It is notable that SA strongly induced a salicylate hydroxylase gene, pointing to a potential SA detoxification mechanism in *Z. tritici*. Salicylate hydroxylases catalyze the degradation of salicylic acid, and similar SA-degrading activities have been described in *Fusarium graminearum*, where fungal genes involved in salicylic acid catabolism contribute to the ability to tolerate and metabolize SA (Rocheleau et al. 2019). In *Z. tritici*, however, the persistent reduction in growth under SA stress suggests that any such detoxification response is insufficient to fully counteract the effects imposed at the concentration tested. We also identified a locus associated with SA response located upstream of a predicted beta-1,3-glucanase gene. Beta-1,3-glucanases are involved in fungal cell wall remodeling and have been linked to pathogenicity in other fungi (Ruiz-Herrera and Ortiz-Castellanos 2019). Beta-1,3-glucanase activity generates beta-1,3-glucans that are critical for maintaining fungal cell wall integrity but also act as potent pathogen-associated molecular patterns (PAMPs) that elicit plant immune responses (Ballou et al. 2016; Liu et al. 2023). Our analyses revealed significant k-mers in the promoter region of the beta-1,3-glucanase gene, suggesting that regulatory variation at this locus may influence its expression. This regulatory variation could shape diverse phenotypic responses to SA stress within the *Z. tritici* population, reflecting an adaptive balance between maintaining cell wall integrity and immune evasion, as shown for other cell-wall degrading enzymes (Rebaque et al. 2025).

Similar to salicylic acid, gibberellic acid reduced fungal growth, but its molecular signature was markedly different. In contrast to the broad transcriptional responses observed under salicylic acid, acidic pH, and prolonged oxidative stress, the GA environment induced only a small number of differentially expressed genes, indicating that its inhibitory effect is accompanied by limited transcriptional effect. This pattern suggests that GA may affect fungal performance through mechanisms that require only subtle changes in gene expression, or through processes acting primarily at the posttranscriptional, posttranslational, or metabolic level. Although some fungal pathogens are known to produce gibberellins that may interfere with host physiology, no gibberellin biosynthetic cluster has been identified in *Z. tritici* (Hassani et al. 2022). In plants, the contribution of GA to defense remains context dependent and is not yet fully resolved (Tanaka et al. 2006; Ding et al. 2013; Gupta and Chakrabarty 2013; Buhrow et al. 2021), although GA is well known to influence plant growth-related traits (Salazar-Cerezo et al. 2018). Consistent with the phenotypic effect of GA in our assays, we identified three loci associated with GA-induced growth phenotypes using GWAS. One early growth-stage locus mapped to a gene encoding an ATP-dependent protease involved in protein turnover and chaperone-related functions, pointing toward proteostasis as a possible component of the GA response.

By contrast, we found that a moderately acidic pH in an otherwise normal growth media enhanced fungal growth. This observation is consistent with reported leaf apoplast pH values, which typically range from 5.0 to 6.0, and suggests that acidic conditions may provide a favorable niche for *Z. tritici* colonization (Felle 2002; Geilfus 2017; Shetty et al. 2008; Gámez-Arjona et al. 2022). At the transcriptional level, acidic pH by itself also triggered one of the strongest responses observed in this study, with extensive differential gene expression and substantial overlap with the late oxidative stress response. In particular, the acidic pH response was enriched for oxidoreductase-associated functions, indicating that low pH is not merely permissive for growth but also elicits active physiological adjustment. Together, these results suggest that *Z. tritici* is well adapted to acidic apoplast-like environments through both enhanced growth and broad transcriptional reprogramming. Consistent with this observation, GWAS identified loci associated with growth under GPL pH 5, including one overlapping a GA-associated locus and another near a predicted E3 ubiquitin ligase-like gene. These results point to a genetic contribution to growth performance in acidic environments and suggest that posttranslational regulation and proteostasis may form part of the adaptive response to apoplast-like conditions.

Oxidative stress showed an altogether different pattern. Although hydrogen peroxide exposure was not the strongest inhibitor of growth, it elicited one of the broadest transcriptomic responses, particularly after prolonged exposure. The early response (15 hpi) to oxidative stress was limited, whereas the 90 hpi treatment induced the second-largest number of environment-specific DEGs and showed the strongest overlap with the acidic pH response, suggesting that the pH 5 and H_2_O_2_ environments activate shared response pathways. These findings are consistent with previous reports that oxidative stress responses in *Z. tritici* depend on exposure conditions and developmental context, including cell density (Francisco et al. 2019). Our results also suggest that oxidative stress adaptation is dynamic, with transcriptional reprogramming becoming more pronounced over time as *Z. tritici* cells adjust to H_2_O_2_ stress. Earlier work proposed that *Z. tritici* can rapidly counteract hydrogen peroxide through multiple peroxidases and catalases encoded in its genome (Amaral et al. 2012). In line with this idea, we detected an over-representation of genes involved in protein binding activities and 55 DEGs with oxidoreductase activity. Despite this pronounced transcriptional response, GWAS did not identify loci significantly associated with oxidative stress-related phenotypes. This absence of significant associations may indicate that oxidative stress tolerance is genetically complex and likely shaped by many loci of small effect, which are difficult to resolve with the present GWAS framework. This interpretation is compatible with earlier QTL mapping that identified two regions on chromosome 8 and chromosome 10 associated with oxidative stress but also linked to growth in other conditions, suggesting a broader role in general stress resilience rather than oxidative stress specificity (Zhong et al. 2021).

## Materials and methods

### Strain collections

This study utilized a diverse collection of 411 *Z. tritici* strains obtained mainly from 21 distinct wheat fields located in 20 countries across major wheat-growing regions between 1981 and 2016 (details in Supplementary Table S1). These strains represent eight distinct genetic clusters, as described by Miñana-Posada et al. (2024), which essentially correspond to geographic continents with some subdivisions in the Americas. The clusters include “Argentina-Uruguay” (31 strains), “Chile” (15 strains), “USA” (129 strains), “Canada” (14 strains), “Europe” (99 strains), “Middle East” (56 strains), “Tunisia” (17 strains), and “Australia” (40 strains). Ten strains, with admixture coefficients below 0.75 (admixed), were not assigned to a genetic cluster and originated from Ukraine (4), Turkey (3), the United States (1), Iran (1), and Ethiopia (1). All the strains were preserved at -80°C in 50% glycerol or anhydrous silica.

### *In vitro* phenotyping in environments simulating host-defense responses

We simulated environments associated with plant defense responses using a modified protocol based on Boixel et al. (2019). Each strain was pre-cultured by transferring 150 µL of glycerol stock into 50 mL of yeast extract peptone dextrose (YPD) medium (10 g/L yeast extract, 20 g/L bactopeptone, 20 g/L dextrose, and 50 mg/L kanamycin). Cultures were incubated at 18°C in the dark with shaking at 120 rpm for seven days. After incubation, cultures were filtered through two layers of gauze to isolate blastospores (yeast-like asexual spores). The filtrate was centrifuged at 3,273 × g for 10 minutes at 4°C, and the resulting spore pellets were resuspended in ∼20 mL of glucose peptone liquid (GPL) medium (14.3 g/L dextrose, 7.1 g/L bactopeptone, 1.4 g/L yeast extract, and 50 mg/L kanamycin).

Spore concentrations were determined by diluting the suspension 1:10 and 1:100 and loading the dilutions onto a hemocytometer (KOVA® GLASSTIC® slide). Spore counts were quantified using a microscope-mounted camera and the Spore Counting v.9 macro in ImageJ (https://github.com/jalassim/SporeCounter). For each strain, spore suspensions were prepared in treatment media with different environments simulating host-defense responses: salicylic acid (SA) was added to a final concentration of 2 mM in GPL medium (adjusted to pH 5 with HCl); gibberellic acid (GA) was also prepared at 2 mM in GPL medium (pH 5); hydrogen peroxide (H_2_O_2_) was used at 1 mM in GPL medium (pH 7); low pH treatment consisted of GPL medium (adjusted to pH 5 with HCl); and control conditions consisted of GPL medium (pH 7).

Each treatment spore suspension had a starting concentration of 2.5 × 10^5^ spores/mL, with 150 µL added to four replicate wells of sterile, flat-bottomed 96-well microtiter plates (Greiner Bio-One, item #655161). Sixteen wells per plate contained only treatment media to serve as absorbance blanks. Plates were sealed with breathable tape (Breathe-Easy®, Diversified Biotech) to allow gas exchange and incubated at 18°C in the dark at 70% humidity for 144 hours.

Fungal growth was monitored by measuring the optical density at 405 nm (OD_405_) twice daily using a plate reader (Spark™ 10M, Tecan). Three technical replicates were performed for each measurement to account for instrument variability. An outlier filtering step was applied based on the standard deviation of the absorbance blank wells for each timepoint to minimize contamination and device errors. To convert OD_405_ values into spore concentrations, calibration curves were generated for each strain using a dilution series ranging from 1 × 10^5^ to 2 × 10^6^ spores/mL. Two interquartile range (IQR) filters were applied to the spore suspension wells to mitigate contaminations and device errors. Linear regression models were fitted using the *lm* function from the *stats* R package and detailed calibration results are provided in Supplementary Table S11. First, an upper-bound IQR filter was applied to each timepoint, plant defense response factor, and replicate, removing 8,406 outliers from 377,208 measurements. A second IQR filter was applied to the median concentration of each strain per condition and timepoint, removing an additional 483 outliers. No more than two replicates per strain were excluded, and the filtering process showed no bias with regard to populations or microtiter plates. Raw absorbance data and are provided in the Zenodo repository: https://doi.org/10.5281/zenodo.19496660.

### Growth variables under environments simulating host-defense responses

To construct individual growth curves, we determined the median spore concentration for each strain, timepoint, and simulated plant defense response environment. These curves were used to extract seven variables to investigate the response of *Z. tritici* strains to host-related stress factors, following a modified version of the method outlined by Miñana-Posada et al. (2024). Logistic regression models, fitted using the *nls* function in R with a self-starting logistic model, were employed to estimate three static growth parameters: growth rate, inflection time, and carrying capacity (i.e., asymptote) for each curve. We then applied local polynomial regressions (*loess* function in R) to interpolate the spore concentrations at 48, 96, and 144 hours. We calculated the empirical area under the curve (eAUC) by averaging the right and left Riemann sums to summarize the overall growth patterns.

Sensitivity to simulated plant defense responses was assessed by calculating the ratio of the eAUC for each factor to the eAUC of its corresponding control. Due to differences in pH between the treatments, for gibberellic acid and salicylic acid, low pH media was used as the control, while for low pH, hydrogen peroxide, and potassium salt, neutral pH media served as the control. The values of each variable, including the sensitivity per strain, can be found in Supplementary Table S11, and summary statistics of each variable per genetic cluster can be found in Supplementary Table S5.

### RNA-sequencing and differential gene expression analysis

The pre-culture of the reference strain IPO323 (Goodwin et al. 2011) was prepared as previously described for plant defense response phenotyping. To assess the transcriptomic response, we prepared 25 mL spore suspensions (2.5 x 10^5^ spores/mL) for each treatment, maintaining the same environmental conditions used for the phenotyping. These suspensions were incubated for 90 hours at 18°C in the dark, with three biological replicates per condition. For hydrogen peroxide, due to its rapid degradation, a spore suspension of the hydrogen peroxide treatment and another spore suspension for the control were prepared and incubated for only 15 hours at 18°C. After incubation, 2 mL of each spore suspension was transferred and centrifuged at approximately 9400 x g for 2 minutes to remove the media. The resulting spore pellets were immediately flash-frozen and stored at -80°C. Frozen samples were sent to Biomarker Technologies GmbH (BMKgene, Münster, Germany) for RNA extraction and sequencing. Sequencing was performed using Illumina NovaSeq X, with paired-end 150 bp reads.

Sequencing adapters and reads shorter than 50 bp were trimmed using Trimmomatic v.0.38 (Bolger et al. 2014) with the following parameters: ILLUMINACLIP:${adapters}:2:30:10 LEADING:15 TRAILING:15 SLIDINGWINDOW:5:15 MINLEN:50. The trimmed reads were aligned to the *Z. tritici* IPO323 transcriptome reference (Lapalu et al. 2023), and transcript abundances were quantified using Kallisto v0.46.1 (Bray et al. 2016) with default parameters for paired-end reads. Read counts were summarized using tximport v1.32.0 (Soneson et al. 2016). Differential gene expression analysis was conducted using the R package *DESeq2* (Love et al. 2014). Principal component analysis (PCA) was performed on the *DESeq2* r-log transformed normalized counts. Genes were considered differentially expressed if the adjusted p-value was < 0.05 and the log2 fold change was > 1.5 (upregulated) or < -1.5 (downregulated)(Love et al. 2014).

### K-mer genome-wide association study

We performed a k-mer-based GWAS, which improves detection power for genome-wide associations (Dutta et al. 2022), following the method of (Voichek and Weigel 2020). This analysis was performed with the same software versions and a similar methodology to that of Miñana-Posada et al. (2024). Sequencing reads were trimmed with Trimmomatic (Bolger et al., 2014) to remove low-quality bases, and 31 bp k-mers were counted to generate a presence/absence table for the strains. A kinship matrix was constructed from the k-mer table and incorporated into GWAS analyses using GEMMA (Zhou and Stephens 2012). K-mers were considered significant if their significance value (*p*-value) reached the 10% family-wise error rate threshold (Voichek and Weigel 2020). Redundant sequences were removed to maintain a unique significant k-mer dataset. Shared significant k-mers between the control environment and the environments simulating host-defense responses were also excluded.

To map significant k-mers, we built a composite reference genome from 19 fully assembled *Z. tritici* genomes (Badet et al. 2020). BLAST searches were performed to identify genomic locations of the significant k-mers, and k-mer loci were defined as significant windows containing more than two k-mers over a maximum of 20 kb. Gene proximity to significant loci was determined using bedtools (Quinlan and Hall 2010), and loci were cross-referenced with orthogroups. Orthogroups were obtained using ProteinOrtho (Lechner et al. 2011). Shared loci were classified based on common orthogroups or k-mers, and each locus was manually inspected to remove duplicates. Gene functions associated with significant loci were explored by referencing the IPO323 annotation (Lapalu et al. 2023) and conducting InterProScan searches (Jones et al. 2014). The overlap with predicted transposable elements (TEs) was evaluated using the latest TE annotations for the 19 *Z. tritici* reference genomes (Baril and Croll 2023). The genomic sequences of the 411 strains used in this study are publicly available, with NCBI Bioproject numbers listed in Supplementary Table S1.

## Code availability

The code for k-mer-based GWAS is available at https://github.com/afeurtey/Kmer_GWAS/tree/main.

## Data availability

The population sequencing data are publicly available from the NCBI Sequence Read Archive with accession numbers shown in Supplementary Table S1. The RNA-seq data generated in this study are available under the Gene Expression Omnibus accession number GSE327738. The raw absorbance data for phenotyping are available in the Zenodo repository: https://doi.org/10.5281/zenodo.19496660.

## Assistance from an AI language model

ChatGPT (OpenAI) was used to provide recommendations to improve grammar and clarity based on input from the authors. All suggestions were carefully reviewed and edited by the authors to ensure the final text accurately conveyed the intended message.

## Author contributions

SMP, AF and CL wrote the manuscript. BAM revised the manuscript. SMP, AF, and CL designed the experiments, analyzed and interpreted the data. SMP acquired the data. SMP, AF, and CL contributed to the conception of the work.

## Acknowledgments

We extend our thanks to ETH Zurich “Hilfsassistierende” student Katja Mühlecker for her assistance with the preparation of *in vitro* phenotyping materials and the initial inoculum of the strains. We are also grateful to MSc student Daniel Osoko for providing the preliminary R scripts used in processing the phenotyping data. Genotyping and growth phenotyping data were generated in collaboration with the Genetic Diversity Centre (GDC), ETH Zurich. This project was supported by funding from ETH Zurich. CL is supported by a SNSF Ambizione grant (project number: PZ00P3_209022).

---

**Supplementary Fig. S1. Pearson’s correlations between the seven growth variables and the sensitivity environments simulating host-defense responses.** The color scale represents the correlation coefficients, while significance levels are denoted by asterisks (*: Benjamini–Hochberg corrected p-value < 0.05; **: Benjamini–Hochberg corrected p-value < 0.01; ***: Benjamini–Hochberg corrected p-value < 0.001).

**Supplementary Fig. S2. Comparisons of the distribution of seven growth variables across environments simulating host-defense responses.** Boxplots displaying per-strain estimates for the seven growth variables across environments simulating host defense responses and in control media. Triangles within each boxplot represent the mean value for each condition. Lowercase letters (a–d) within each growth variable panel denote significant differences among environments, as determined by Tukey’s post-hoc pairwise comparisons based on linear mixed-effects models.

**Supplementary Fig. S3. Sensitivity distribution across environments simulating host-defense responses and genetic clusters**. (A) Individual graphs showing boxplots of sensitivities for each treatment across genetic clusters. (B) Individual graphs displaying boxplots of sensitivities for each genetic cluster across treatments. Triangles within each boxplot indicate the mean sensitivity for each environment and genetic cluster. Lowercase letters (a–d) denote significant differences between treatments or genetic clusters, as determined by Tukey’s post-hoc pairwise comparisons based on a linear mixed-effects model. If no lowercase letters are displayed, no significant differences were detected.

**Supplementary Fig. S4. Empirical area under the curve (eAUC) distribution across environments simulating host-defense responses and genetic clusters**. Boxplots showing the eAUC for each genetic cluster across environments. Triangles within each boxplot indicate the mean eAUC for each environment and genetic cluster. Lowercase letters (a–d) denote significant differences between genetic clusters, as determined by Tukey’s post-hoc pairwise comparisons based on a linear mixed-effects model. If no lowercase letters are displayed, no significant differences were detected.

**Supplementary Fig. S5. Growth rate distributions across environments simulating host-defense responses and genetic clusters**. A) Individual graphs showing boxplots of growth rates for each environment across genetic clusters. B) Individual graphs displaying boxplots of growth rates for each genetic cluster across environments. Triangles within each boxplot indicate the mean sensitivity for each environment and genetic cluster. Lowercase letters (a–d) denote significant differences between environments or genetic clusters, as determined by Tukey’s post-hoc pairwise comparisons based on a linear mixed-effects model. If no lowercase letters are displayed, no significant differences were detected.

**Supplementary Fig. S6. Limited overlap of differentially expressed genes (DEGs) across environments simulating host-defense responses.** A) Pairwise similarity of DEG sets between environments, quantified using the Jaccard index (intersection/union). Values are shown for comparisons among 90 hpi SA (vs GPL pH 5), 90 hpi H_2_O_2_ (vs GPL pH 7), 90 hpi GPL pH 5 (vs GPL pH 7), 15 hpi H_2_O_2_ (vs GPL pH 7), and 90 hpi GA (vs GPL pH 5). (B) UpSet plot summarizing the overlap of DEGs across environments. Horizontal bars indicate the total number of DEGs per environment, while vertical bars show the size of each intersection. The dot–line matrix below indicates which environments contribute to each intersection. **(C)** Expression profiles of the nine DEGs shared across four of the five environments (SA pH 5, GA pH 5, GPL pH 5, 90 hpi H_2_O_2_, and 15 hpi H_2_O_2_; each relative to its matched control). Points indicate log2 fold-change in each environment; colors denote direction of regulation (green, up-regulated; purple, down-regulated).

**Supplementary Fig. S7. Functional composition of environment-specific differentially expressed genes (DEGs).** Heatmaps showing the number of environment-specific DEGs assigned to broad functional categories for each environment comparison. The left panel summarizes environment-specific down-regulated DEGs and the right panel summarizes environment-specific up-regulated DEGs. Columns correspond to pairwise contrasts (GA pH 5 vs GPL pH 5; GPL pH 5 vs GPL pH 7; 15 hpi H_2_O_2_ pH 7 vs 15 hpi GPL pH 7; 90 hpi H_2_O_2_ pH 7 vs 90 hpi GPL pH 7; SA pH 5 vs GPL pH 5), and rows indicate functional categories. Cell values give the number of DEGs in each category, with shading reflecting counts. Asterisks denote categories significantly enriched among environment-specific DEGs for a given treatment relative to the background set of functionally annotated IPO323 genes (Fisher’s exact test with FDR correction; * FDR < 0.05, ** FDR < 0.01, *** FDR < 0.001).

**Supplementary Fig. S8. Top up-regulated differentially expressed genes (DEGs) across environments simulating host defense responses.** Heatmap summarizing log2 fold-change (log2FC) values for the most strongly up-regulated genes identified in each differential expression comparison: GPL pH 5 vs GPL pH 7, GA pH 5 vs GPL pH 5, 15 hpi H_2_O_2_ pH 7 vs 15 hpi GPL pH 7, 90 hpi H_2_O_2_pH 7 vs 90 hpi GPL pH 7, and SA pH 5 vs GPL pH 5. Rows correspond to gene models (gene IDs; brief functional annotations shown where available). Genes are grouped by the environment in which they rank among the top induced transcripts (horizontal separators). Color indicates the magnitude of induction (green to purple, increasing log2FC; scale at right). Cells are shown for the corresponding comparison; genes not significantly induced in a given comparison are displayed near baseline.

**Supplementary Table S1.** Details for each strain, including the strain name, alternative names used in other publications, sampling year, genetic cluster classification by Miñana-Posada et al. (2024), and the Bioproject accession ID associated with the publicly available genomic sequencing data.

**Supplementary Table S2.** Pearson’s correlation coefficients and the Benjamini–Hochberg corrected p-values between the seven growth variables and the sensitivity across environments simulating host defense responses.

**Supplementary Table S3.** ANOVA of each of the LMERs per the seven growth variables and the sensitivity to environments simulating host defense responses.

**Supplementary Table S4.** Post hoc Tukey pairwise comparisons between environments simulating host defense responses for each growth variable and sensitivity.

**Supplementary Table S5.** Summary statistics of the seven growth variables and the sensitivity per environment simulating host defense responses.

**Supplementary Table S6.** Post hoc Tukey pairwise comparisons between environments simulating host defense responses and genetic cluster for eAUC, growth rate, and sensitivity.

**Supplementary Table S7.** Differential gene expression results for comparisons between environments. Genes were classified as “Upregulated” if the adjusted *p*-value was < 0.05 and the log2 fold change exceeded 1.5, or as “Downregulated” if the log2 fold change was less than -1.5. Genes not meeting these criteria were categorized as “Not Significant.”

**Supplementary Table S8.** Significantly associated k-mers with the seven phenotypic variables and sensitivity under environments simulating host defense responses.

**Supplementary Table S9.** Genomic loci significantly associated with phenotypes under environments simulating host defense responses.

**Supplementary Table S10.** Orthogroup analysis from ProteinOrtho for the 19 *Z. tritici* references.

**Supplementary Table S11.** Growth variables and sensitivity estimates for each strain of the 411 *Z. tritici* strains per environment simulating host defense responses and the control environment.

