## Supplementary Fig. S for "Contrasting transcriptional responses and genetic determinants underlie *Zymoseptoria tritici* adaptation mechanisms to simulated host defense environments"

Miñana-Posada et al.

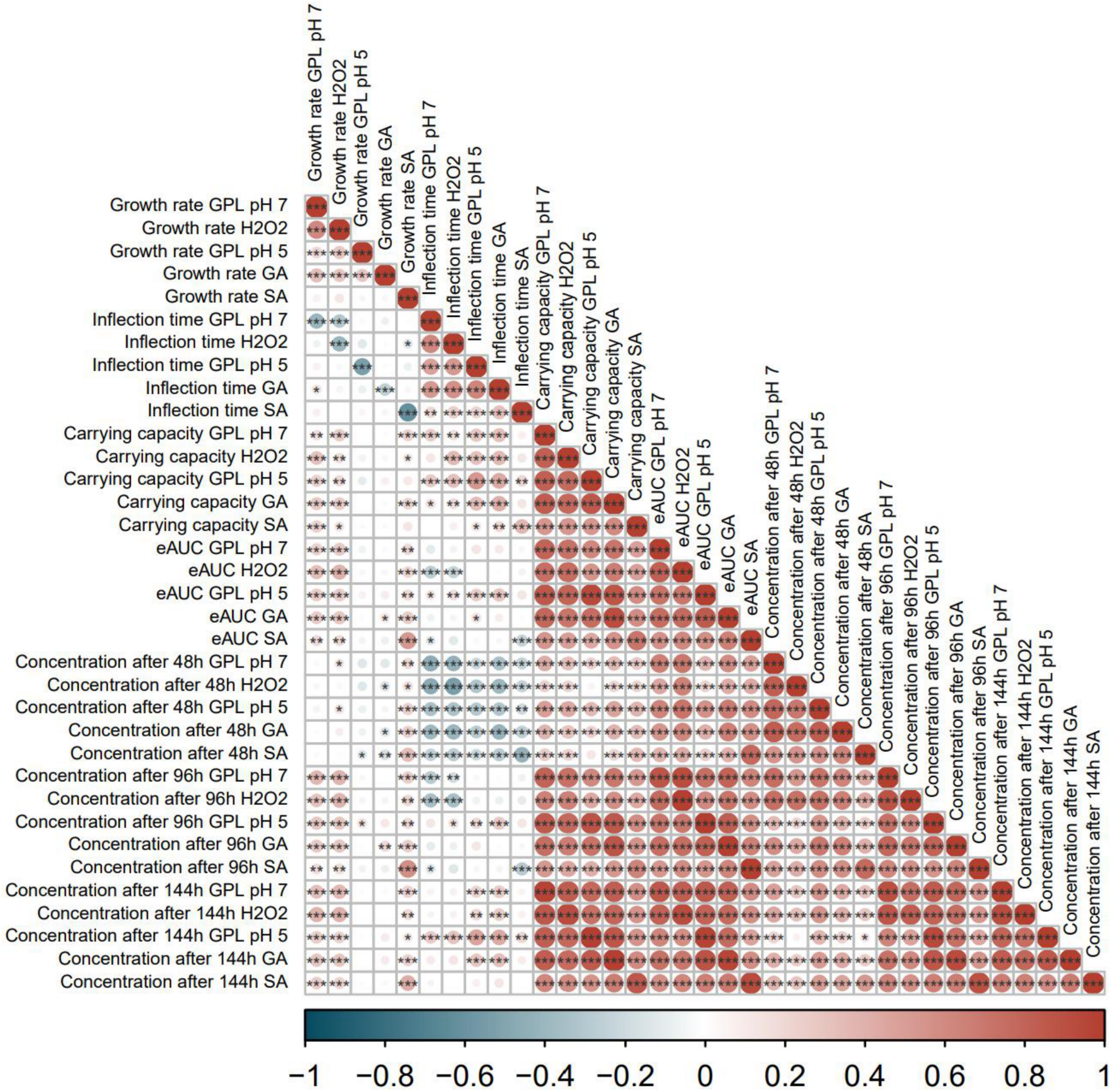


**Supplementary Fig. S1. Pearson's correlations between the seven growth variables and the sensitivity environments simulating host-defense responses**. The color scale represents the correlation coefficients, while significance levels are denoted by asterisks (*: Benjamini–Hochberg corrected p-value < 0.05; **: Benjamini–Hochberg corrected p-value < 0.01; ***: Benjamini–Hochberg corrected p-value < 0.001).

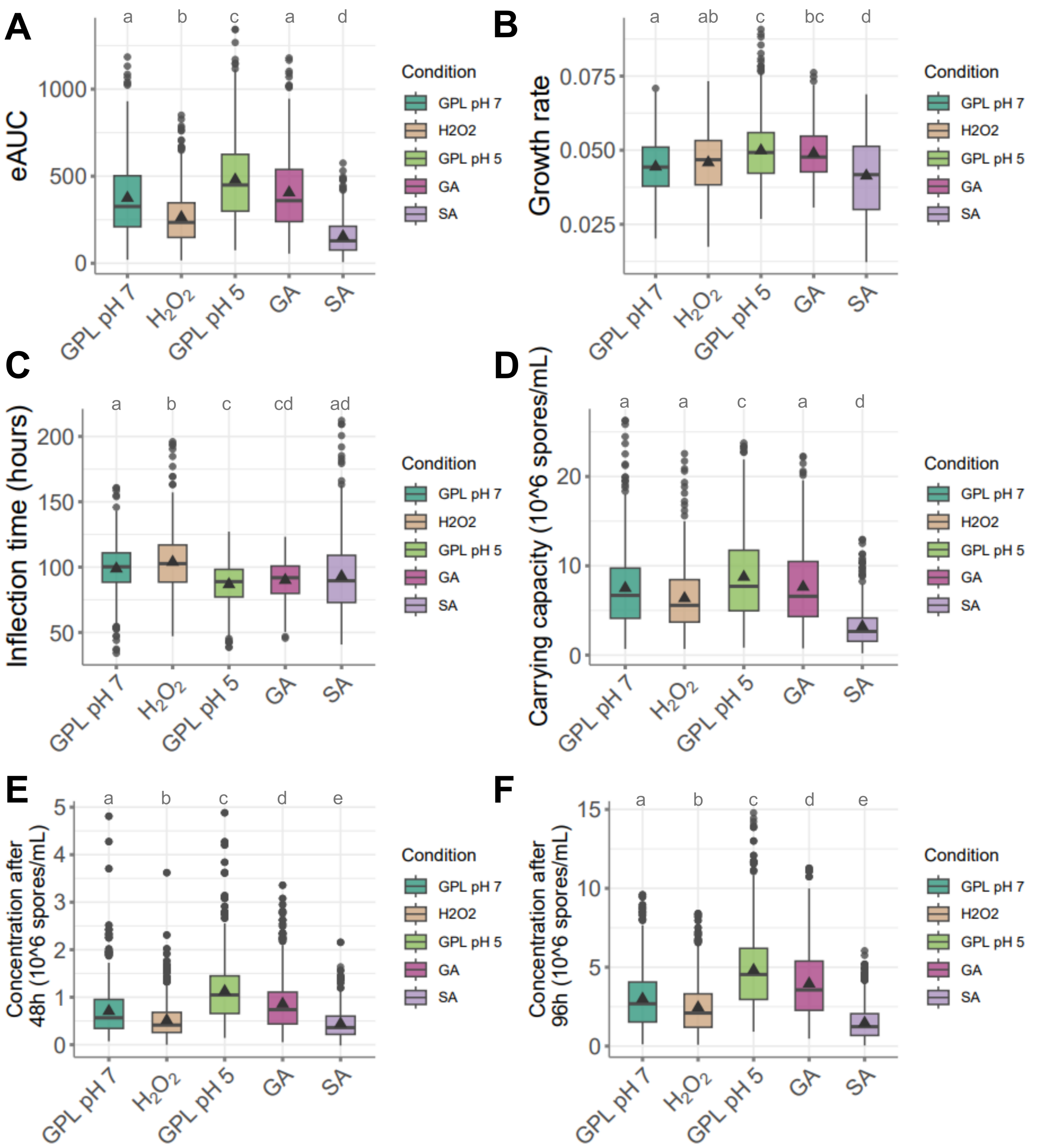


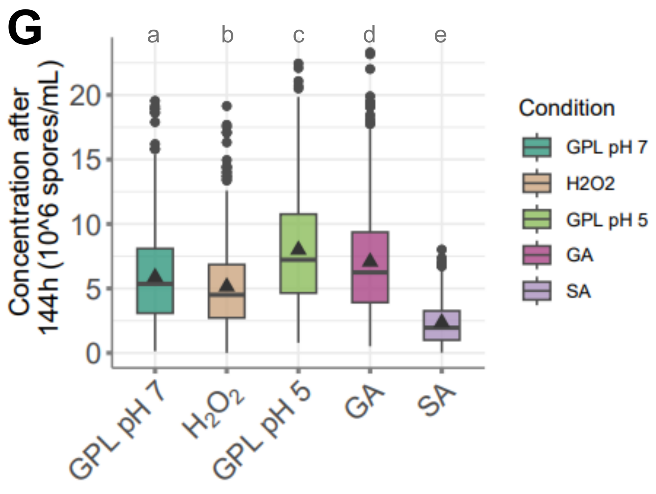


**Supplementary Fig. S2. Comparisons of the distribution of seven growth variables across environments simulating host-defense responses.** Boxplots displaying per-strain estimates for the seven growth variables across environments simulating host defense responses and in control media. Triangles within each boxplot represent the mean value for each condition. Lowercase letters (a–d) within each growth variable panel denote significant differences among environments, as determined by Tukey's post-hoc pairwise comparisons based on linear mixed-effects models.

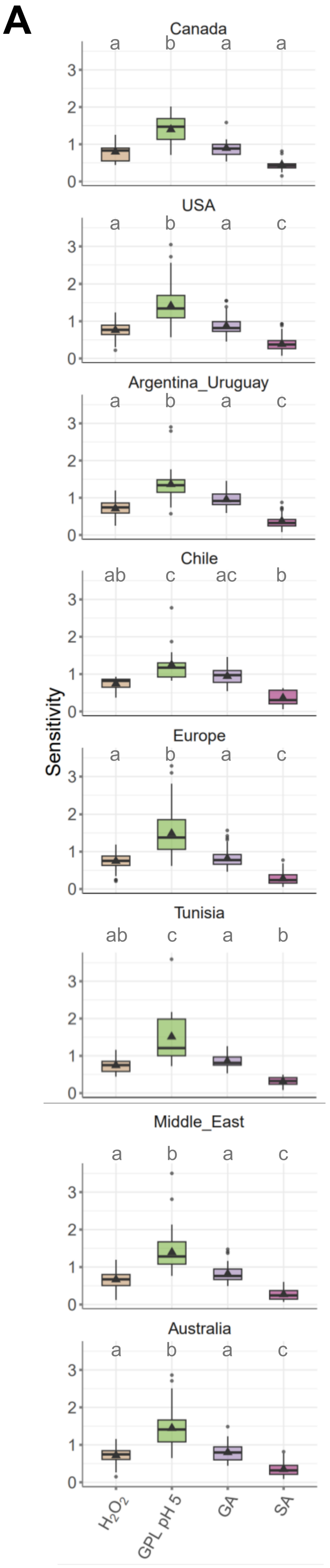


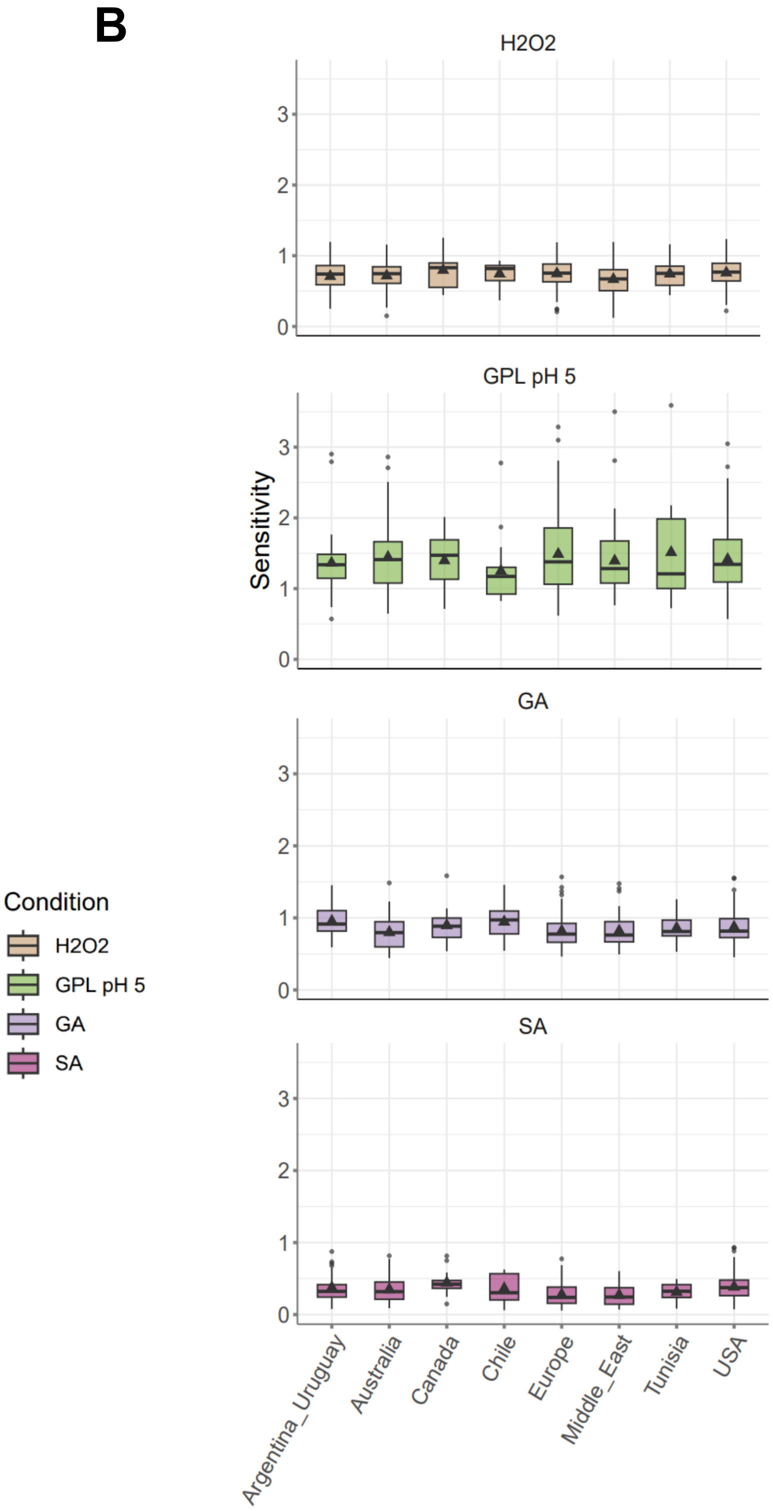


**Supplementary Fig. S3. Sensitivity distribution across environments simulating host defense responses and genetic clusters.** (A) Individual graphs showing boxplots of sensitivities for each treatment across genetic clusters. (B) Individual graphs displaying boxplots of sensitivities for each genetic cluster across treatments. Triangles within each boxplot indicate the mean sensitivity for each environment and genetic cluster. Lowercase letters (a–d) denote significant differences between treatments or genetic clusters, as determined by Tukey's post-hoc pairwise comparisons based on a linear mixed-effects model. If no lowercase letters are displayed, no significant differences were detected.

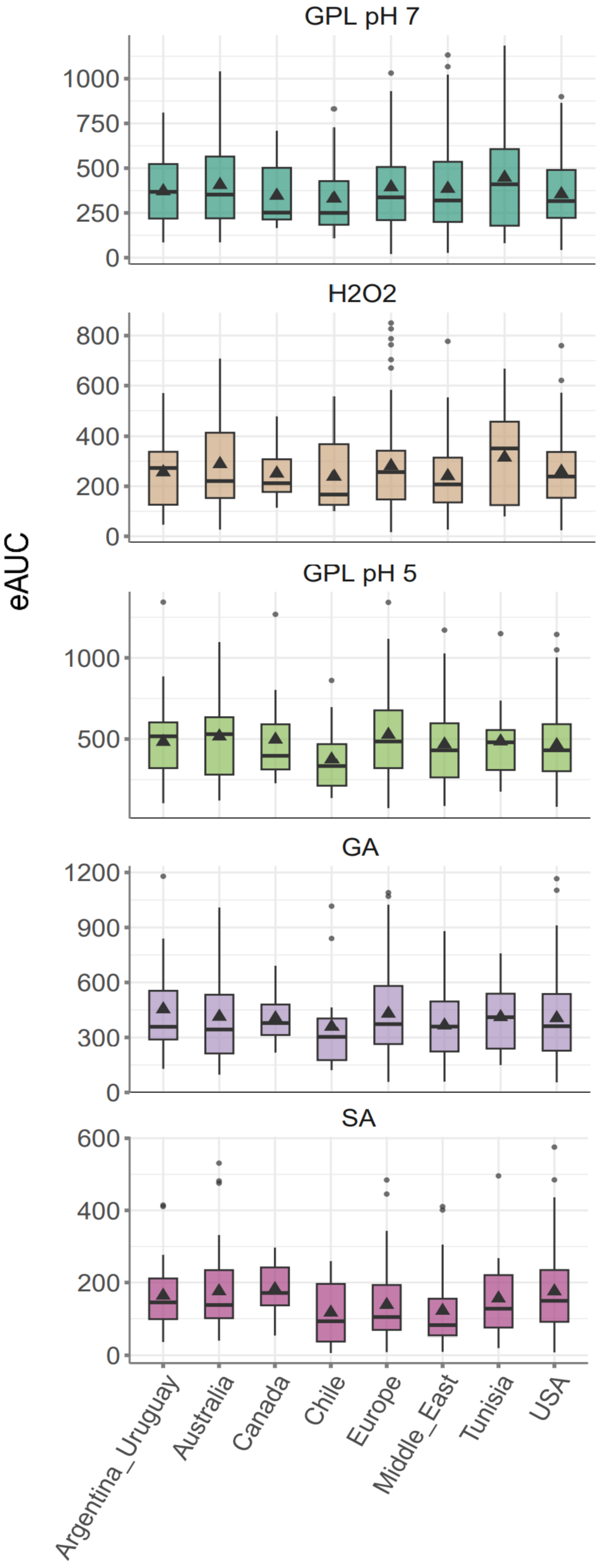


**Supplementary Fig. S4. Empirical area under the curve (eAUC) distribution across environments simulating host defense responses and genetic clusters.** Boxplots showing the eAUC for each genetic cluster across environments. Triangles within each boxplot indicate the mean eAUC for each environment and genetic cluster. Lowercase letters (a–d) denote significant differences between genetic clusters, as determined by Tukey's post-hoc pairwise comparisons based on a linear mixed-effects model. If no lowercase letters are displayed, no significant differences were detected.

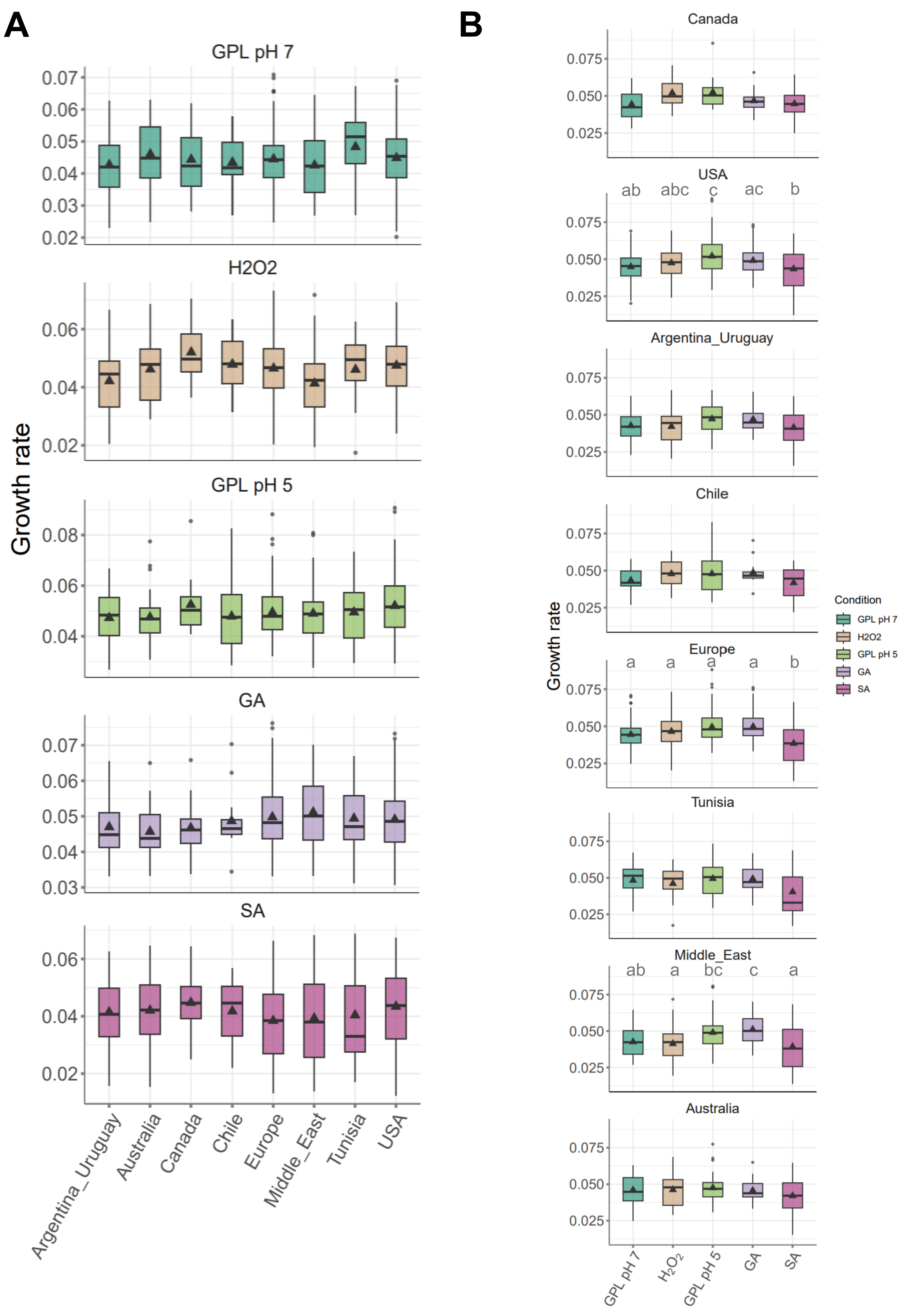


**Supplementary Fig. S5. Growth rate distributions across environments simulating host defense responses and genetic clusters.** A) Individual graphs showing boxplots of growth rates for each environment across genetic clusters. B) Individual graphs displaying boxplots of growth rates for each genetic cluster across environments. Triangles within each boxplot indicate the mean sensitivity for each environment and genetic cluster. Lowercase letters (a–d) denote significant differences between environments or genetic clusters, as determined by Tukey's post-hoc pairwise comparisons based on a linear mixed-effects model. If no lowercase letters are displayed, no significant differences were detected.

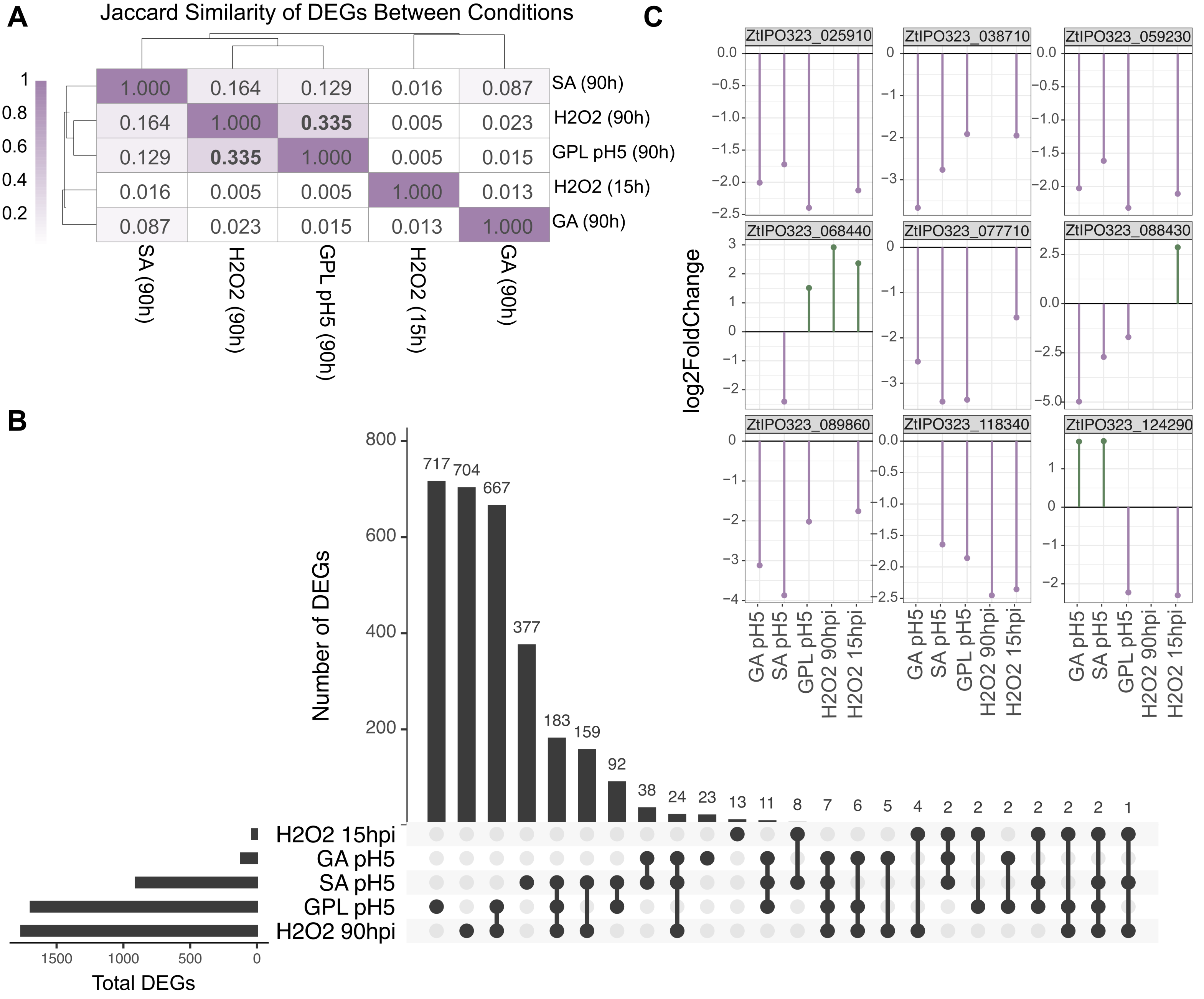


**Supplementary Fig. S6. Limited overlap of differentially expressed genes (DEGs) across environments simulating host defense responses.** A) Pairwise similarity of DEG sets between environments, quantified using the Jaccard index (intersection/union). Values are shown for comparisons among 90 hpi SA (vs GPL pH 5), 90 hpi H2O2 (vs GPL pH 7), 90 hpi GPL pH 5 (vs GPL pH 7), 15 hpi H2O2 (vs GPL pH 7), and 90 hpi GA (vs GPL pH 5). (B) UpSet plot summarizing the overlap of DEGs across environments. Horizontal bars indicate the total number of DEGs per environment, while vertical bars show the size of each intersection. The dot–line matrix below indicates which environments contribute to each intersection. (C) Expression profiles of the nine DEGs shared across four of the five environments (SA pH 5, GA pH 5, GPL pH 5, 90 hpi H2O2, and 15 hpi H2O2; each relative to its matched control). Points indicate log2 fold-change in each environment; colors denote direction of regulation (green, up-regulated; purple, down-regulated).

**Condition-specific down-regulated DEGs**

| | GA pH5 vs GPL pH5 | GPL pH5 vs GPL pH7 | 15hpi $H_2O_2$ pH7 vs 15hpi GPL pH7l | 90hpi H2O2 pH7 vs 90hpi GPL pH7 | SA ph5 vs GPL pH5 |
|---|---|---|---|---|---|
| Transmembrane transport | 1 | 22 | 0 | 25 | 12 |
| Oxidoreductase activity | 0 | 14 | 0 | 13 | 16 |
| ATP binding | 0 | 13 | 0 | 10 | 6 |
| Transporter activity | 1 | 19 | 0 | 15 | 8 |
| Protein binding | 0 | 18 | 0 | 6** | 15 |
| Membrane component | 0 | 19 | 0 | 24 | 17 |
| Zinc ion binding | 0 | 10 | 0 | 8 | 7 |
| Proteolysis | 1 | 6 | 0 | 5 | 5 |
| Heme binding | 0 | 4 | 0 | 9 | 5 |
| Carb. metabolism | 0 | 12 | 0 | 4 | 3 |

**Condition-specific up-regulated DEGs**

| | GA pH5 vs GPL pH5 | GPL pH5 vs GPL pH7 | 15hpi $H_2O_2$ pH7 vs 15hpi GPL pH7 | 90hpi H2O2 pH7 vs 90hpi GPL pH7 | SA ph5 vs GPL pH5 |
|---|---|---|---|---|---|
| Transmembrane transport | 1 | 10 | 2 | 9 | 8 |
| Oxidoreductase activity | 0 | 32*** | 1 | 14 | 6 |
| ATP binding | 1 | 11 | 1 | 11 | 2 |
| Transporter activity | 1 | 6 | 2 | 6 | 6 |
| Protein binding | 0 | 7 | 1 | 10 | 2 |
| Membrane component | 2 | 10 | 1 | 7 | 6 |
| Zinc ion binding | 0 | 2 | 0 | 2 | 3 |
| Proteolysis | 0 | 7 | 3 | 4 | 1 |
| Heme binding | 0 | 4 | 1 | 6 | 1 |
| Carb. metabolism | 0 | 6 | 0 | 7 | 1 |

**Supplementary Fig. S7. Functional composition of environment-specific differentially expressed genes (DEGs).** Heatmaps showing the number of environment-specific DEGs assigned to broad functional categories for each environment comparison. The left panel summarizes environment-specific down-regulated DEGs and the right panel summarizes environment-specific up-regulated DEGs. Columns correspond to pairwise contrasts (GA pH 5 vs GPL pH 5; GPL pH 5 vs GPL pH 7; 15 hpi H2O2 pH 7 vs 15 hpi GPL pH 7; 90 hpi H2O2 pH 7 vs 90 hpi GPL pH 7; SA pH 5 vs GPL pH 5), and rows indicate functional categories. Cell values give the number of DEGs in each category, with shading reflecting counts. Asterisks denote categories significantly enriched among environment-specific DEGs for a given treatment relative to the background set of functionally annotated IPO323 genes (Fisher's exact test with FDR correction; * FDR < 0.05, ** FDR < 0.01, *** FDR < 0.001).

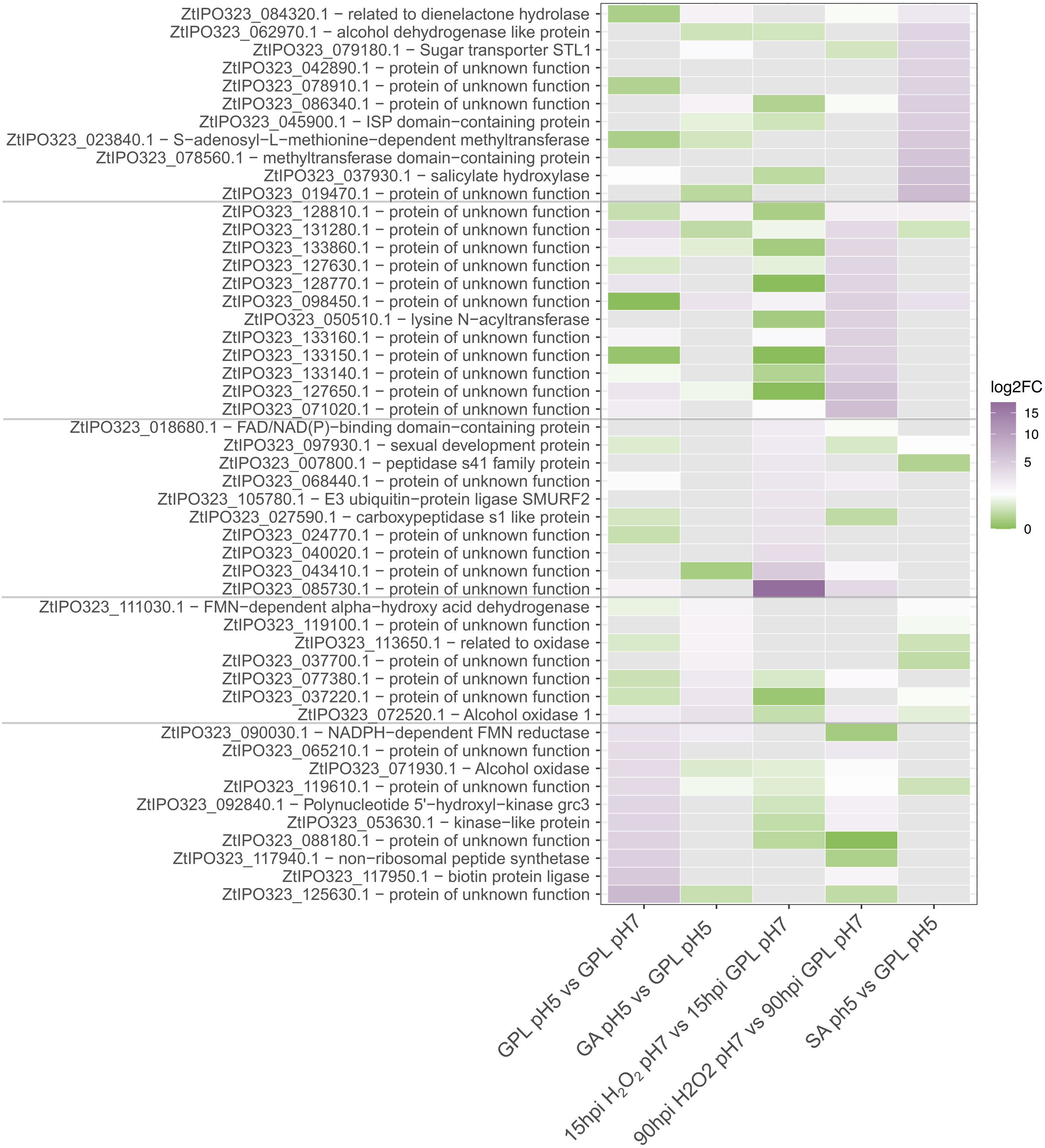


**Supplementary Fig. S8. Top up-regulated differentially expressed genes (DEGs) across environments simulating host defense responses.** Heatmap summarizing log2 fold-change (log2FC) values for the most strongly up-regulated genes identified in each differential expression comparison: GPL pH 5 vs GPL pH 7, GA pH 5 vs GPL pH 5, 15 hpi H2O2 pH 7 vs 15 hpi GPL pH 7, 90 hpi H2O2pH 7 vs 90 hpi GPL pH 7, and SA pH 5 vs GPL pH 5. Rows correspond to gene models (gene IDs; brief functional annotations shown where available). Genes are grouped by the environment in which they rank among the top induced transcripts (horizontal separators). Color indicates the magnitude of induction (green to purple, increasing log2FC; scale at right). Cells are shown for the corresponding comparison; genes not significantly induced in a given comparison are displayed near baseline.
